# Reward learning deficits in Parkinson’s disease depend on depression

**DOI:** 10.1101/069062

**Authors:** Monique H.M. Timmer, Guillaume Sescousse, Marieke E. van der Schaaf, Rianne A.J. Esselink, Roshan Cools

**Affiliations:** Donders Institute for Brain, Cognition and Behaviour, Centre for Cognitive Neuroimaging, Radboud University, 6500 HB Nijmegen, the Netherlands; Radboud university medical center, Department of Neurology and Parkinson Center Nijmegen (ParC), 6500 HB Nijmegen, the Netherlands; Radboud university medical center, Department of Psychiatry, 6500 HB Nijmegen, the Netherlands

## Abstract

**Background:** Depression is one of the most common and debilitating non-motor symptoms of Parkinson’s disease (PD). The neurocognitive mechanisms underlying depression in PD are unclear and treatment is often suboptimal.

**Methods:** We investigated the role of striatal dopamine in reversal learning from reward and punishment by combining a controlled medication withdrawal procedure with functional magnetic resonance imaging (fMRI) in 22 non-depressed PD patients and 19 PD patients with past or present PD-related depression.

**Results:** PD patients with a PD-related depression (history) exhibited impaired reward versus punishment reversal learning as well as reduced reward versus punishment-related BOLD signal in the striatum (putamen) compared with non-depressed PD patients. No effects of dopaminergic medication were observed.

**Conclusions:** The present findings demonstrate that impairments in reversal learning from reward versus punishment and associated reward-related striatal signalling depend on the presence of (a history of) depression in PD.

## Introduction

Patients with Parkinson’s disease (PD) experience not only motor symptoms, such as bradykinesia and rigidity, but also non-motor symptoms, such as cognitive/affective deficits, psychosis and depression. Depression is one of the most frequently observed and debilitating non-motor symptoms of PD with a prevalence of 30-40% (Reijnders et al., 2008). Despite its high prevalence and impact, the neurobiological and -cognitive mechanisms underlying depression in PD are unclear and accordingly, treatment is often suboptimal.

Depression has been associated with an imbalance in the impact of reward and/or punishment on learning, behaviour and cognition (Clark et al., 2009, Eshel and Roiser, 2010, Roiser et al., 2012, Der-Avakian and Markou, 2012, Whitton et al., 2015, Treadway and Zald, 2013). For example, patients with depression have been shown to exhibit both enhanced impact of punishment as well as reduced impact of reward on learning (Murphy et al., 2003, Robinson et al., 2011, Taylor Tavares et al., 2008). Notably, negative affective biases are also observed in individuals at risk for depression in several cognitive domains, including learning (Forbes et al., 2007, Robinson et al., 2010a, Roiser et al., 2012), putatively representing a vulnerability factor (Eshel and Roiser, 2010). In this study, we asked whether similar biases in learning from reward versus punishment contribute to depression in PD.

This question is particularly relevant given extensive evidence that PD is accompanied by dopamine-dependent changes in the balance between reward-versus punishment-based learning, a facet of cognition that critically relies upon dopaminergic prediction error coding in the striatum (Schultz and Dickinson, 2000). Multiple studies have shown that dopaminergic medication in PD patients reduces punishment-based learning, while, if anything, enhances reward-based learning (Smittenaar et al., 2012, Cools et al., 2006, Frank et al., 2004, Bodi et al., 2009, Palminteri et al., 2009, Rutledge et al., 2009, Moustafa et al., 2008). According to current modeling work, these drug effects reflect dopamine-induced shifts in the balance between activity in the direct and indirect pathways of the basal ganglia (Frank, 2005). Despite consistent medication effects, discrepancy exists between studies with regard to the pattern of performance on such tasks of PD patients OFF medication. While some studies report performance to be unaltered in the OFF state compared with controls (Cools et al., 2006, Rutledge et al., 2009, Smittenaar et al., 2012, Moustafa et al., 2008), other studies report that reward-based learning is actually impaired relative to punishment (Bodi et al., 2009, Kobza et al., 2012, Frank et al., 2004, Palminteri et al., 2009). The pattern of impaired reward versus punishment learning in PD patients OFF medication resembles that described above in depressed individuals (non-PD) (Clark et al., 2009, Eshel and Roiser, 2010) and concurs generally with suggestions that striatal dopamine depletion contributes to depression in PD. For instance, nuclear neuroimaging studies revealed that depression PD is accompanied by decreased dopamine transporter binding, especially in ventral striatal regions, compared with non-depressed patients (Remy et al., 2005, Vriend et al., 2013, Weintraub et al., 2005). Functional MRI studies in depressed individuals (non-PD) have shown attenuated ventral striatal functioning across various tasks (Epstein et al., 2006, Forbes et al., 2009, Pizzagalli et al., 2009), including reward-based learning (Robinson et al., 2011). Based on this evidence, we hypothesized that the presence of impaired reward versus punishment learning in PD patients OFF medication might depend on the presence of (a history of) depression and associated ventral striatal dysfunction.

Specifically, we predicted that depressed PD patients, OFF medication, would exhibit a greater imbalance between learning from reward versus punishment and greater abnormalities in ventral striatal BOLD signal than non-depressed PD patients. Moreover, this negative learning bias and associated ventral striatal dysfunction in PD-related depression would be remedied by dopaminergic medication. Thus, we expected dopaminergic medication to normalize reward versus punishment learning and associated ventral striatal BOLD signal in depressed patients, while impairing punishment versus reward learning and associated ventral striatal BOLD signal in non-depressed patients (cf(Cools et al., 2006)).

To test these hypotheses, we investigated effects of dopaminergic medication withdrawal in PD patients with and without (a history of) PD-related depression, using pharmacological fMRI and a well-established reversal learning paradigm. This paradigm was designed specifically to disentangle reward-from punishment-based reversal learning and previous fMRI work with this paradigm has shown that both unexpected reward and unexpected punishment elicit a prediction error signal in the striatum (Robinson et al., 2010b). Moreover, this paradigm has been shown to be sensitive to dopaminergic manipulation in healthy volunteers and PD patients (Cools et al., 2009, van der Schaaf et al., 2014, Janssen et al., 2015, Cools et al., 2006) as well as depression (non-PD)(Robinson et al., 2011). Here we build on this prior work to advance our understanding of the neurochemical and neurocognitive mechanisms of depression in PD.

## Materials and Methods

### Participants and general procedure

Twenty-four depressed and 23 non-depressed PD patients were recruited. Data from 5 depressed patients and 1 non-depressed patient were excluded from the analysis. Two depressed patients failed to complete the study, leading to incomplete datasets. One depressed patient turned out to be claustrophobic and was not able to perform the task inside the MRI scanner. Three PD patients (2 depressed and 1 non-depressed) were excluded because of outlying behaviour (mean error rates across the task as a whole >3SD from the group mean). Therefore, results are based on datasets from 19 depressed patients and 22 non-depressed patients. This is an appropriate sample size for cognitive fMRI studies with a between-group design (Thirion et al., 2007).

This study was part of a larger project investigating the neurobiological mechanisms of depression in PD. All participants gave informed consent as approved by the local research ethics committee (CMO region Arnhem - Nijmegen, The Netherlands, nr. 2012/43) and were compensated for participation. Patients were recruited from the Parkinson Centre at the Radboud university medical center, Nijmegen, the Netherlands, and were diagnosed with idiopathic PD according to the UK Brain Bank criteria by a neurologist specialized in movement disorders (Prof. B.R. Bloem, Dr. R.A. Esselink, Dr. B. Post). All patients used dopaminergic medication (non-depressed group: levodopa n=10, dopamine receptor agonists n=2, combination of both n=10; depressed group: levodopa n=14, dopamine receptor agonists n=2, combination of both n=3). Patient groups were matched for the amounts of daily dopaminergic medication use (levodopa equivalent dose (Esselink et al., 2004), t(39)=1.22, p=0.23) as well as the amounts of daily dopamine receptor agonist use (t(39)=1.47, p=0.15). Six depressed patients used antidepressants (Paroxetine n=2, Escitalopram n=1, Citalopram n=1 and Nortriptyline n=2). All patients were on stable medication regimes during the course of the study, except for one patient who used Duloxetine, a serotonin/noradrenalin reuptake inhibitor, for 4 weeks between the two testing days. The drug was prescribed to treat pain and discontinued 4 weeks before the second testing day.

Exclusion criteria were clinical dementia (Mini Mental State Examination <24 (Folstein et al., 1975)), psychiatric disorders other than depression, neurological co-morbidity and hallucinations. Patients were assigned to the depressed group if they met the DSM-IV criteria, based on structured psychiatric interviews conducted during an intake session (MINI-plus, (Sheehan et al., 1998)), for a major (n=5) or minor (n=12) depressive episode, dysthymic disorder (n=1) or adjustment disorder with depressed mood (n=1) within five years before PD diagnosis up until now. Five patients suffered from depressive symptoms during the course of the study. The other patients were diagnosed with past PD-related depression. The incidence of depression is significantly higher within the five years preceding PD diagnosis and is therefore likely related to PD pathology (Shiba et al., 2000). This criterion was chosen, because we expected PD patients with past or present PD-related depression to exhibit similar negative learning biases, reflecting an underlying vulnerability. The groups were matched for age, gender, IQ (Dutch version of the National Adult Reading Test (Schmand et al., 1991)), disease severity (Unified Parkinson Disease Rating Scale part III (Goetz and Stebbins, 2004)) and amounts of dopaminergic medication (Levodopa Equivalent Dose (Esselink et al., 2004)) (Table1).

**Table 1.** Patient characteristics. Values represent number of patients or mean (SD). NART = National Adult Reading Test, MMSE = Mini Mental State Examination, UPDRS = Unified Parkinson Disease Rating Scale, LED = Levodopa Equivalent Dose, BDI = Beck Depression Inventory, averaged across the ON and OFF session in patients.

|  | Depressed<br>n = 19 | Non-depressed<br>n = 22 |
| --- | --- | --- |
| Gender, men | 12 | 14 |
| Age, years | 58.4 (5.3) | 61.1 (7.6) |
| NART-IQ | 96.0 (11.5) | 97.8 (15.0) |
| MMSE | 28.5 (1.3) | 28.6 (1.2) |
| Handedness, right | 16 | 18 |
| Response hand, right | 5 | 14 |
| UPDRS-III (OFF) | 23.3 (9.4) | 21.9 (6.8) |
| LED (mg/day) | 527 (240) | 626 (277) |
| LED agonists (mg/day) | 55 (114) | 110 (127) |
| BDI (averaged) | 8.7 (5.0) | 4.3 (2.3) |
| First session ON | 9 | 13 |
| Days between sessions | 24.1 (28.6) | 21.5 (20.2) |

Patients were assessed on two occasions - once while taking their normal dopaminergic medication (ON) and once after withdrawal from their dopaminergic medication for at least 18 hours (24 hours for controlled-release dopamine receptor agonists) (OFF). Antidepressants were not withdrawn, enabling us to assess specifically dopaminergic drug effects. The order of OFF and ON sessions was counterbalanced in each group (Table1). During testing days, participants performed the task described below. Furthermore, current depressive symptoms were measured using the Beck Depression Inventory (BDI). Testing days always started in the morning between 8:30-10:30 am.

### Task

We used a deterministic reversal learning paradigm (Figure 1) similar to that used in previous studies (Cools et al., 2006, Robinson et al., 2011, van der Schaaf et al., 2014). The task was presented on a screen visible via a mirror attached on the head coil in the MRI scanner. On each trial, participants were shown 2 simultaneously presented vertically adjacent stimuli, 1 scene and 1 face. One of these stimuli was associated with reward, the other with punishment. By trial and error, subjects had to learn these deterministic stimulus-outcome associations. Unlike classical instrumental reversal learning paradigms, subjects did not choose between stimuli, but had to predict whether the highlighted stimulus was associated with reward or punishment. Subjects indicated their prediction by pressing the reward or punishment button with their least affected hand. Response mappings were counterbalanced across subjects. Stimuli were presented until a response was made, after which the actual outcome was shown. If subjects did not respond in time, a “Too late” message was presented. Stimulus-outcome contingencies reversed after 4-6 consecutive correct predictions. Reversals were signalled by either an unexpected reward (presented after a highlighted stimulus that was previously associated with punishment) or an unexpected punishment (presented after a highlighted stimulus that was previously associated with reward). Unexpected outcomes were only presented after a correct prediction was made according to the current contingency ruling-out the possibility of reversal anticipation. Moreover, participants were informed that reversal anticipation was not possible within the structure of this task. The same stimulus was always highlighted again on the first trial after an unexpected outcome to ensure that a contingency reversal would always be paired with a reversal in motor response. Patients were familiarized with the task during the intake session and performed a practice block on each testing day.

**Figure 1.**
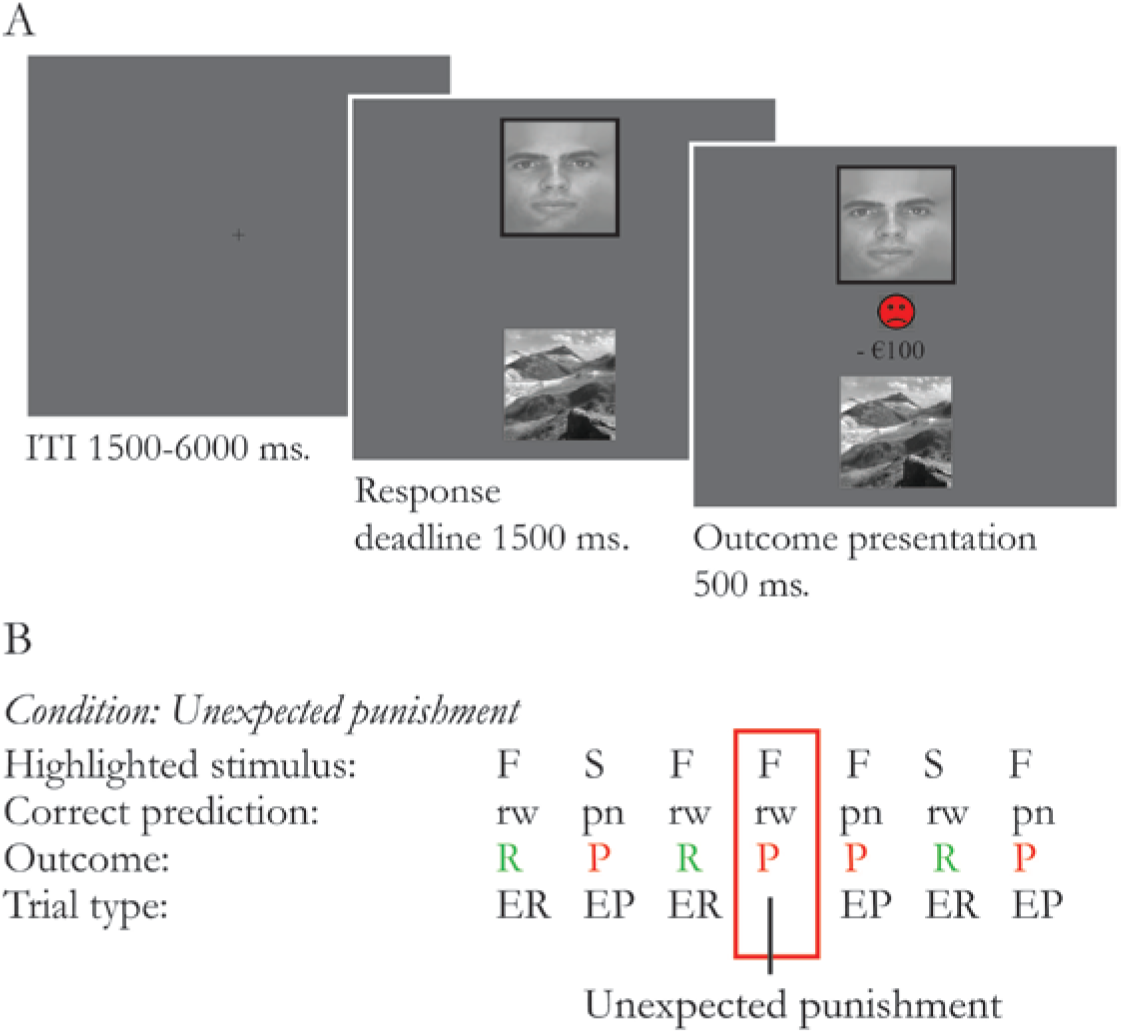
Task overview. Two stimuli (a face and a scene) were simultaneously presented. One of the stimuli was highlighted with a black border. Participants were asked to predict if the highlighted stimulus was followed by reward (green happy smiley and “+€100” sign) or punishment (red sad smiley and “-€100” sign). Following the participants’ prediction, the actual outcome was presented (100% deterministic). B Example sequence of trials. In this example the face stimulus was associated with expected reward (ER) and the scene stimulus was associated with expected punishment (EP). After a series of 4 to 6 consecutive correct responses, the stimulus-outcome associations reversed, signaled by either unexpected reward or unexpected punishment.

On each testing day, subjects completed 2 experimental blocks of 230 trials. Each experimental block contained a short break of 30s. The number of reversals depended on task performance and thus varied across participants. The average number of reversal trials for reward and punishment was 29(±6) and 29(±5), respectively, across groups and did not differ between groups or drug sessions.

### Behavioural analysis

Error rates and reaction times were analyzed with a mixed ANOVA with GROUP as a between-subject factor and REVERSAL (reversal, non-reversal), VALENCE (reward, punishment) and DRUG (OFF and ON medication) as within-subject factors. Errors were defined as misses or incorrect predictions. Errors on reversal trials were defined as errors (i.e. incorrect predictions) on the trial immediately following an unexpected outcome. All other trials were defined as non-reversal trials, including trials that were followed by an unexpected outcome. Note that unexpected outcomes only followed a correct prediction according to the current contingency. Thus, errors on trials that were followed by an unexpected outcome could not occur within the structure of this task. Error rates were arcsine transformed (2*arcsin(√ *x*)) as is appropriate when variance is proportionate to the mean (Howell 1997).

### Image acquisition and analysis

A Siemens TIM-Trio 3-T MRI scanner with a 32-channel head-coil was used to acquire structural and functional MRI images. Functional images were acquired using a multi-echo echoplanar imaging sequence (38 axial slices, ascending slice acquisition order, voxel size = 3.3×3.3×2.5mm, matrix=64×64, repetition time (TR)=2.32s, echo time (TE)=9.0/19.3/30.0/40.0ms, flip angle=90°). Multi-echo images were acquired in order to benefit from reduced susceptibility artifacts at low echo times (Poser et al., 2006). The structural image was acquired using a T1-weighted MP-RAGE sequence (192 slices, voxel size=1.0×1.0×1.0mm, matrix=256×256, TR=2.3s, TE=3.03s, flip-angle=8°).

Images were preprocessed and analyzed using SPM8 (Welcome Department of Cognitive Neurology, London). Images were realigned to the first volume using data from the shortest TE to estimate realignment parameters. After realignment, a weighted summation was performed to combine all four TEs into a single dataset (Poser et al., 2006). To this aim, thirty “resting-state” images, acquired before the start of the actual experiment, were used to estimate BOLD contrast-to-noise ratio maps for each TE. These maps were then used to calculate an optimal voxel-wise weighting between the four echoes using in-house software, maximizing the contribution of each echo according to its contrast-to-noise ratio. Combined images were checked for spiking artefacts, slice-time corrected to the middle slice, coregistered to the structural image, normalized to the standard Montreal Neurological Institute template, re-sampled into 2.5×2.5×2.5mm isotropic voxels and smoothed with an isotropic Gaussian kernel of 8mm full-width at half-maximum.

A first-level general linear model (GLM) was estimated that incorporated separate regressors for each possible outcome (modelled as event at time of outcome presentation, convolved with a canonical hemodynamic response function): unexpected punishment, unexpected reward, correctly predicted expected punishment, correctly predicted expected reward, incorrectly predicted expected outcomes and miss trials. An additional epoch regressor modelled the 30s break within each experimental block. Twenty-nine noise regressors were added to the GLM: 24 motion regressors (6 derived from the realignment procedure, their first derivatives (n=6) and those squared (n=12)), 3 parameters to model global intensity changes (the time series of the mean signal from white matter, cerebral spinal fluid and out-of brain segments) and 2 regressors to control for BOLD signal changes related to (changes in) tremor amplitude; an electromyography (EMG) amplitude regressor and its first derivative both convolved with a canonical hemodynamic response function (Helmich et al., 2011). Time series were high-pass filtered (cut-off 128sec) to remove low-frequency signals and an AR(1) model was applied to adjust for serial correlations. The 2 experimental blocks from 1 session were modelled within the same GLM. Preprocessing and estimation of the GLM was performed separately for each drug session.

Individual contrast maps were generated at the first level for each drug session. The main contrast of interest was [unexpected reward – unexpected punishment]. We calculated individual ‘drug-difference maps’ (OFF-ON) and ‘drug-average maps’ ((OFF+ON)/2). These contrast maps were taken to a second-level random-effects analysis. To compare drug-effects between depressed and non-depressed patients, we submitted individual ‘drug-difference maps’ to a second level two-sample T-test. To assess the main effect of drug, we submitted individual ‘drug-difference maps’ to a second level one-sample T-test and to assess the main effect of group, we submitted individual ‘drug-average maps’ to a second level two-sample T-test. Response hand was added as a covariate of no-interest to control for differences in response hand between groups (Table 1).

Statistical inference was performed at the voxel level using a family-wise error (FWE) corrected threshold of p<0.05 within an a priori defined small-volume of interest corresponding to the striatum (bilateral nucleus accumbens, caudate nucleus and putamen, based on the Automated Anatomical Labeling atlas (p_sv_fwe_)). For additional whole brain analyses, statistical inference was performed at the cluster level using an FWE-corrected threshold of p<0.05 (p_wb_fwe_). Marsbar software was used to extract mean parameter estimates to assess brain-behaviour correlations and for illustration purposes.

## Results

### Patient and disease characteristics

As expected, patient groups differed significantly in depressive symptoms (BDI averaged across the 2 drug sessions, *F_(1,39)_=*=13.22, *p*=0.001, ηp^2^=0.25) (Table 1). Note however that BDI scores of the depressed patient group still fell within the normal range (mean 8.7±5.0) as a result of our inclusion criteria.

### Behavioural results

Task performance in general was very good (Table 2). Comparison of error rates in non-depressed PD patients and PD patients with a PD-related depression (history) revealed a significant three-way interaction of REVERSAL (reversal, non-reversal), VALENCE (reward, punishment) and GROUP (*F_(1,39)_* =4.17, *p*=0.048, ηp^2^=0.10). Breakdown of this interaction revealed a significant REVERSAL*VALENCE interaction in depressed patients (*F_(1,39)_* =8.55, *p*=0.009, ηp^2^=0.32), but not in non-depressed patients (*p*=0.4). The significant interaction in depressed patients was driven by an effect of VALENCE on reversal trials (*F_(18)_* =4.86, *p*=0.041, ηp^2^=0.21). Depressed patients made more errors on reward reversal trials compared with punishment reversal trials. There was also a significant effect of REVERSAL on reward trials (F(_18_) =5.12, *p*=0.036, ηp^2^=0.22), indicating that depressed patients made more errors on reward reversal trials compared with reward non-reversal trials. There was no effect of VALENCE on non-reversal trials (*p*=0.6). There were no other significant interactions with GROUP or DRUG and no significant main effects of GROUP, DRUG, REVERSAL or VALENCE (Figure 2). There were no session order effects. Analyses of reaction times are reported in the supplementary materials.

**Figure 2.**
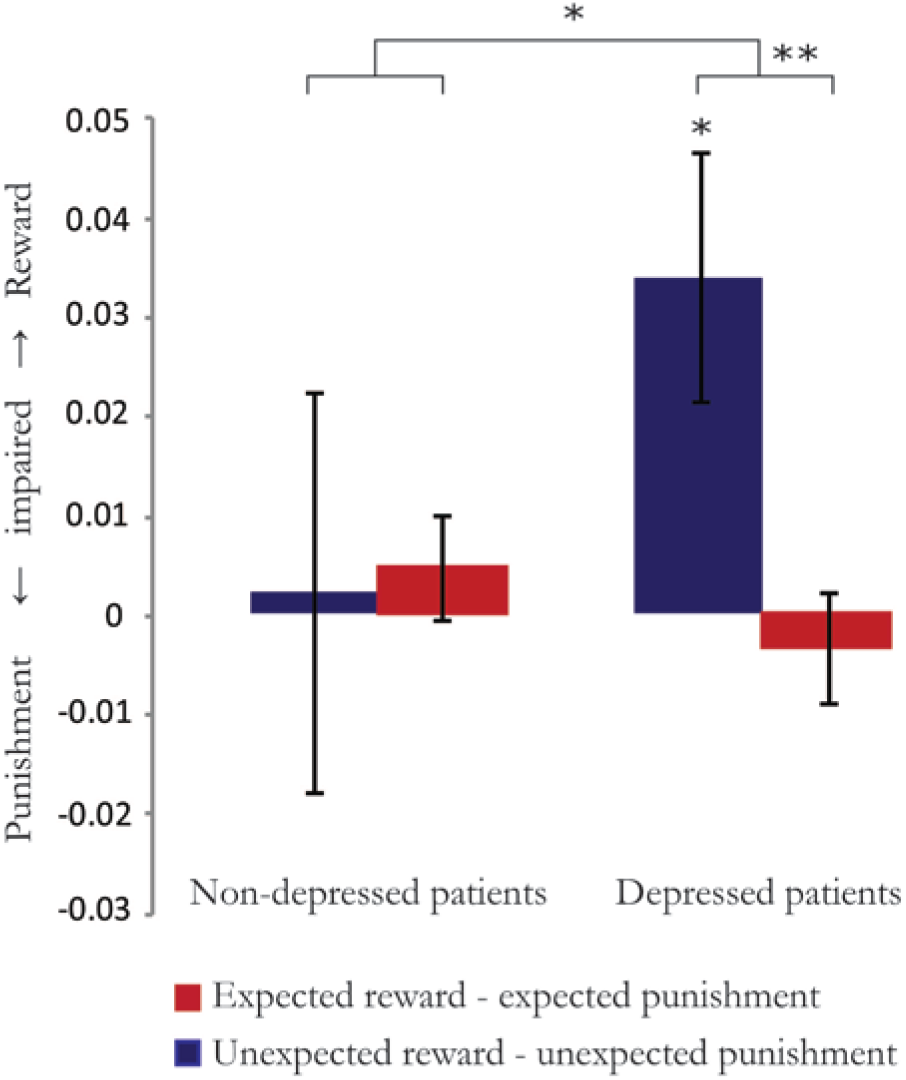
Error rates. Error rates on reversal trials [unexpected reward - unexpected punishment] (in blue) and non-reversal trials [expected reward - expected punishment] (in red) as a function of group (depressed and non-depressed PD patients). Error bars represent standard errors of the mean. \**p*<0.05, ** *p*<0.01, \*\*\**p*<0.001

**Table 2.** Error rates. Median error rate (interquartile range) per group and drug session. EP = expected punishment, UP = unexpected punishment, ER = expected reward, UR = unexpected reward.

| Trial type | Depressed patients | Non-depressed patients |
| --- | --- | --- |
| EP OFF | 0.07 (0.13) | 0.07 (0.10) |
| UP OFF | 0.06 (0.17) | 0.07 (0.10) |
| ER OFF | 0.08 (0.10) | 0.09 (0.08) |
| UR OFF | 0.12 (0.21) | 0.06 (0.12) |
| EP ON | 0.07 (0.08) | 0.10 (0.11) |
| UP ON | 0.07 (0.15) | 0.09 (0.10) |
| ER ON | 0.07 (0.06) | 0.10 (0.06) |
| UR ON | 0.14 (0.14) | 0.09 (0.12) |

### Dopamine receptor agonists

In contrast to previous studies (cf(Cools et al., 2006, Frank et al., 2004)), we did not observe valence-specific effects of dopaminergic medication on reversal learning. Because previous literature suggests that valence-specific drug effects might be driven by patients on dopamine receptor agonists (Cools et al., 2006), we performed a supplementary analysis including dopamine receptor agonist use (AGONIST) as additional between-subject factor. This analysis revealed no significant interactions with GROUP, DRUG or AGONIST as factor(s) and no significant main effects of GROUP, DRUG or AGONIST.

### Antidepressants

In the depressed group 6 patients used antidepressants. A supplementary analysis with ANTIDEPRESSANT use as an additional between-subject factor revealed the same three-way interaction as described above (REVERSAL*VALENCE*GROUP, *F_(1,38)_*=4.30, *p*=0.045, ηp^2^=0.10), but no significant interactions with ANTIDEPRESSANT as a between subject factor.

### Imaging results

We were primarily interested in valence-specific striatal BOLD signal changes during unexpected outcomes in depressed versus non-depressed PD patients. Supplementary analyses on outcome-general reversal-related brain signal changes and on valence-specific brain signal changes during expected outcomes can be found in the supplementary materials. First, given the behavioural results, we assessed group differences using a two-sample T-test on individual ‘drug-average maps’ contrasting unexpected reward and punishment. This analysis revealed a significant group effect on striatal BOLD signal elicited by unexpected reward versus unexpected punishment (small volume analysis: right putamen, x=30, y=-14, z=12, *T*=5.05, p_sv_fwe_=0.008) (Figure 3A,B). Decomposition of this interaction revealed that unexpected reward induced significantly greater increases in striatal BOLD signal than unexpected punishment in non-depressed patients (right putamen, x=30, y=-14, z=12, *T*=5.11, p_sv_fwe_=0.037; left putamen, x=-28, y=-4, z=12, *T*=4.95, p_sv_fwe_=0.049). This effect was not observed in depressed patients (Figure 3A,B). There were no differences in striatal BOLD signal elicited by either unexpected reward or unexpected punishment (contrasted against baseline) between depressed and non-depressed patients, indicating that the observed difference in valence-specific striatal BOLD signal during unexpected outcomes was driven by the difference between reward versus punishment. The effect was restricted to the striatum: There were no other effects elsewhere in the brain as revealed by whole brain analysis. There was also no GROUP*DRUG interaction nor a main effect of DRUG on striatal BOLD signal elicited by reward versus punishment reversal trials.

**Figure 3.**
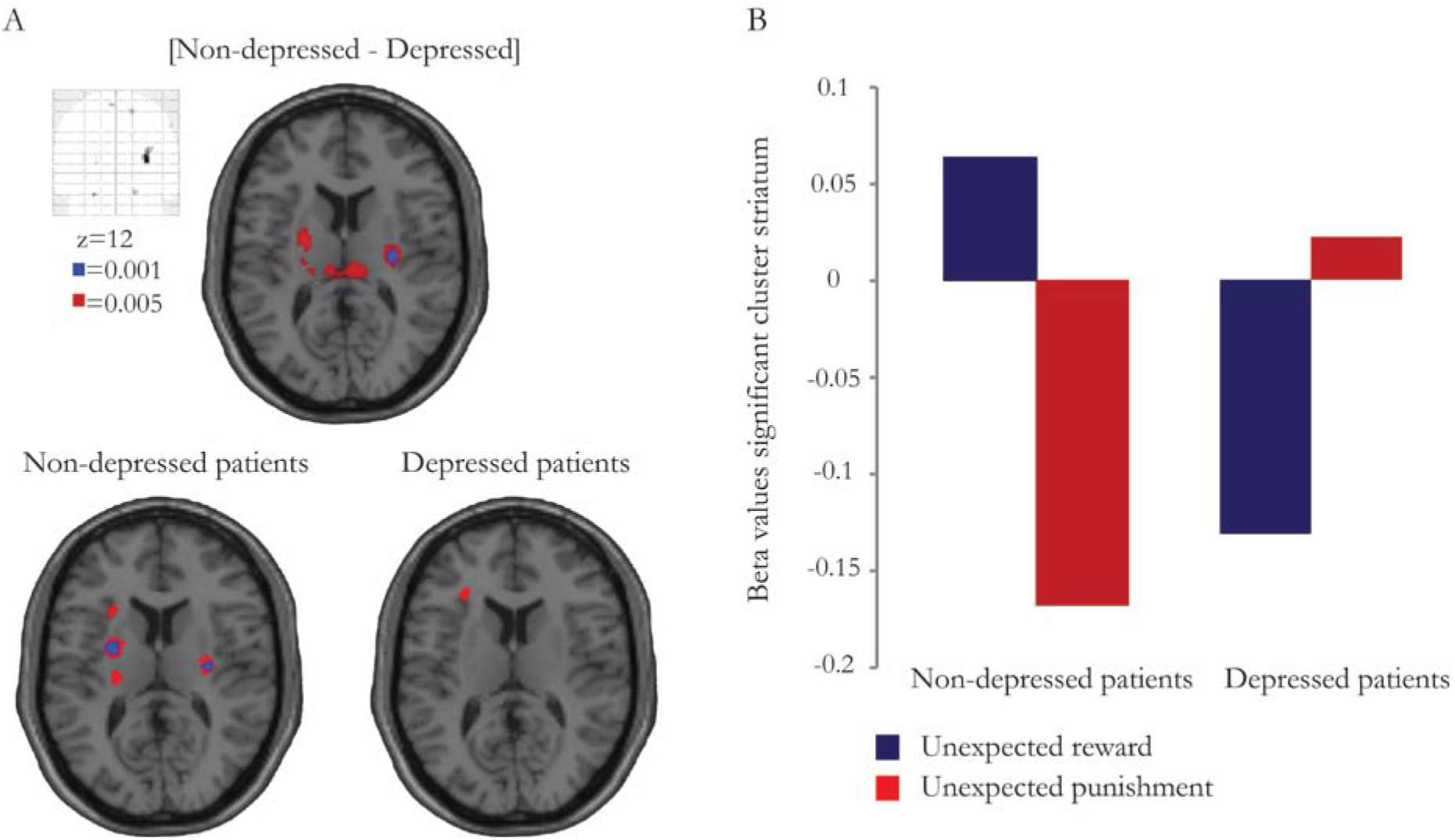
BOLD signal during reward-versus punishment-based reversal learning. Valence-specific BOLD signal in the striatum during unexpected outcomes for the contrast [non-depressed - depressed patients] and for both groups separately (non-depressed patients and depressed patients). Data presented at *p*<0.001 uncorrected (blue) and at *p*<0.005 uncorrected (red). Effects are significant within our small volume of interest (p_sv_fwe_<0.05). B Beta values for unexpected reward and unexpected punishment per group extracted from the significant striatal cluster (right putamen, x=30, y=-14, z=12; 9 voxels) for the contrast [non-depressed – depressed patients].

In patients (across both groups), there was no significant correlation between BDI scores and impairments in valence-specific reversal learning (*r_(41)_*=0.060, *p*=0.71). The correlation between BDI scores and valence-specific BOLD signal changes in the striatum during unexpected outcomes just failed to reach significance (*r_(41)_*=-0.303, *p*=0.054) (individual beta values extracted from the significant striatal cluster; 9 voxels).

## Discussion

In line with our hypothesis, we demonstrate that PD-related depression (history) is accompanied by impaired reward versus punishment reversal learning and attenuated striatal signaling. Specifically, we show impaired reward (versus punishment) reversal learning and reduced striatal BOLD signal in PD patients with a PD-related depression (history) compared with non-depressed patients. Whereas unexpected reward induced significantly greater increases in striatal BOLD signal than unexpected punishment in non-depressed patients, this was not observed in depressed patients.

In depression, impaired reward processing and attenuated striatal function has been shown previously across multiple facets of cognition (Epstein et al., 2006, Forbes et al., 2009, Pizzagalli et al., 2009, Steele et al., 2007). In fact, the present effect concurs directly with a finding from previous work, using the exact same paradigm, showing reduced reward-based reversal learning and reduced striatal signalling (albeit in a slightly more anterior region) in depressed individuals (non-PD)(Robinson et al., 2011). However, this is the first study demonstrating impaired reward (versus punishment) reversal learning and attenuated striatal function in patients with PD and PD-related depression compared with non-depressed patients, thereby providing evidence that abnormal striatal signaling, the key region affected by PD, also contributes to depression-specific cognitive deficits in PD.

There is discrepancy in extant literature with respect to the integrity of reward and/or punishment learning in PD patients OFF medication. Some studies have reported OFF state performance to be unaltered compared with controls (Cools et al., 2006, Moustafa et al., 2008, Rutledge et al., 2009, Smittenaar et al., 2012), whereas other studies have revealed impaired reward relative to punishment learning/performance (Bodi et al., 2009, Kobza et al., 2012, Frank et al., 2004, Palminteri et al., 2009). The current data suggest that these discrepancies might reflect differences in the inclusion of patients with or without a PD-related depression (history). As such, our observations demonstrate the importance of taking into account depression history in PD patients when investigating reward (versus punishment) learning.

The present study demonstrates attenuated striatal responses in depressed PD patients in a rather posterior striatal region. This contrasts with some previous studies in depressed individuals (non-PD) showing blunted striatal responses in more anterior striatal regions (Epstein et al., 2006, Robinson et al., 2011, Steele et al., 2007). Of course this discrepancy might reflect the effect of PD in our study. Critically, a similar posterior striatal locus of reward versus punishment prediction error coding has been previously shown using this exact same paradigm in healthy subjects (Robinson et al., 2010b). In this study, unexpected reward elicited significantly greater BOLD signal increases than unexpected punishment in a posterior striatal region. This was argued to reflect recruitment of instrumental mechanisms in the context of reward (Robinson et al., 2010b). Work with experimental animals as well as functional MRI studies in human revealed that anterior striatal regions are implicated in the prediction of salient stimuli during Pavlovian learning, whereas posterior striatal regions are implicated in outcome-guided action selection and the instrumental control of behaviour (Yin et al., 2006, Montague et al., 1996, O’Doherty et al., 2004). In the current study, contrasting effects of unexpected reward and punishment prediction error coding on posterior striatal BOLD signal were observed in non-depressed, but not in PD patients with a PD-related depression (history). This could reflect the inability of depressed patients to recruit reward-guided instrumental actions. Studies in depressed individuals (non-PD) using classical reinforcement learning paradigms indeed revealed a lack of behavioural adjustment in the face of reinforcing (rewarding) stimuli in depressed (Henriques et al., 1994, Pizzagalli et al., 2005) consistent with diminished reward-guided instrumental control of behaviour in depression. In the current study disentangling Pavlovian from instrumental mechanisms was not possible, leaving this question open for further investigation.

In contrast to our hypothesis, and contrary to previous studies that have consistently reported valence-specific dopaminergic drug effects on (reversal) learning (Bodi et al., 2009, Cools et al., 2006, Frank et al., 2004, Palminteri et al., 2009, Moustafa et al., 2008), we did not observe valence-specific drug effects. We are puzzled by this lack of effect and provide two possible accounts. First, valence-specific drug effects on (reversal) learning have been shown primarily with dopamine receptor agonists and antagonists (Bodi et al., 2009, Cools et al., 2006, Cools et al., 2009, Janssen et al., 2015, Moustafa et al., 2008, van der Schaaf et al., 2014). In contrast to previous studies, in our sample only less than half of the patients used dopamine receptor agonists (17/41). Moreover, most patients in our sample (15/17) used controlled-release dopamine receptor agonists for which one might argue that the withdrawal period was too short. However, the behavioural pattern (across both patient groups) observed in the current study was more akin to that seen in previous studies when patients were in an OFF rather than an ON state, suggesting that the effects of controlled-release dopamine receptor agonists on valence-specific (reversal) learning might not be comparable to those of regular dopamine receptor agonists. A second, not mutually exclusive possibility is that our failure to observe the predicted medication effect might reflect a ceiling effect: In the present study patients performed extremely well, and much better than did the patients in our previous study (Cools et al., 2006). The median error rate OFF (across patients groups) for unexpected punishment was 0.06 and 0.08 for unexpected reward in the current study, while it was 0.12 for unexpected punishment and 0.20 for unexpected reward in our previous study (Cools et al., 2006). Thus, it is possible that there was insufficient dynamic range for any medication-induced improvement in valence-specific learning to surface.

In the depressed patient group, 6 patients used antidepressants; 4 used a serotonin reuptake inhibitor and 2 used a tricyclic antidepressant. It is well known that other neurotransmitters than dopamine, such as serotonin, can influence reward versus punishment learning (Cools et al., 2008, Robinson et al., 2012). A possible effect of antidepressant medication (or an interaction with dopaminergic medication) on task performance cannot be ruled out, although this is unlikely given that prior work has shown that manipulations of central serotonin repeatedly elicited qualitatively different behavioural changes than did various dopamine manipulations (Cools et al., 2008, Robinson et al., 2012). Indeed, a supplementary analysis did not reveal any effect of antidepressant medication (nor an interaction with dopaminergic medication) on task performance.

A potential caveat of the present study is the heterogeneous sample of PD patients with PD-related depression, that included patients with current as well as past PD-related depression. Although the sample sizes of both patients groups (n=19 and n=22) were large enough for a cognitive fMRI study with a between-group design (Thirion et al., 2007), we lacked power for comparison of PD patients with current (n=5) versus past (n=14) PD-related depression. Negative (learning) biases have been shown in never-depressed individuals at risk for depression (Robinson et al., 2010a, Forbes et al., 2007). Moreover, outside the domain of learning it has been shown that negative affective biases can persist after remission of a depressive episode (see for review Roiser et al., 2012). However, the hypothesis that negative learning biases diminish (or persist) with remission of a depressive episode has never been investigated. The present results should therefore be validated in a follow-up study that includes a larger group of PD patients with PD-related depression enabling comparison of PD patients with past and current PD-related depressive symptoms.

To summarize, this is the first study demonstrating that PD-related depression (history) is accompanied by impaired reward (versus punishment) reversal learning and associated striatal signalling. These results demonstrate that attenuated striatal signalling might underlie reward learning deficits in PD patients and shows that striatal reward learning deficits in PD depend on the presence of a depression (history).

## Acknowledgements

We would like to thank all participants for their cooperation in the study. Furthermore, we are grateful to Dr. Rick Helmich and Michiel Dirkx for their help with analyzing the data.

## Ethical standards

The authors assert that all procedures contributing to this work comply with the ethical standards of the relevant national and institutional committees on human experimentation and with the Helsinki Declaration of 1975, as revised in 2008.

